# Immunomodulatory Effects of Selenium on T-Cell-Mediated Responses in Thyroid Eye Disease

**DOI:** 10.64898/2026.04.08.717334

**Authors:** Hemlata Bisnauthsing, Wai Kit Chu

## Abstract

**Background:** Thyroid Eye Disease (TED) is an autoimmune orbital disorder driven by pathogenic T-cell subsets, including T-helper 1 (Th1) and follicular helper T (Tfh) cells, which sustain orbital inflammation and thyroid-stimulating immunoglobulin (TSI) production. Selenium supplementation has demonstrated clinical benefit in mild TED, yet its immunological mechanisms remain poorly defined.

**Methods:** A murine TED model was established in female BALB/c mice via TSHR plasmid immunisation. Animals maintained on a low-selenium diet (0.07 ppm) received sodium selenite supplementation at 0.2 mg/kg/day. Orbital pathology was assessed by immunohistochemistry, H&E and Masson’s Trichrome staining. T-cell subset abundance was quantified by flow cytometry, and serum T4, TRAb, and IL-21 levels were measured by ELISA. In vitro dose-response experiments examined the effects of selenium on Tfh cell viability, IL-21 production, apoptosis, and ferroptosis.

**Results:** Selenium supplementation reduced CD3⁺ T-cell orbital infiltration, collagen fibrosis, and serum T4 and TRAb levels in TSHR-immunised mice. Flow cytometry revealed significant reductions in Tfh and Th1 cell abundance, with Th17 cells unaffected. Serum IL-21 and B-cell abundance were also markedly reduced in vivo. In vitro, selenium exhibited a biphasic, dose-dependent effect on Tfh cells: low concentrations maintained viability and IL-21 production, while higher concentrations induced ferroptosis and apoptosis.

**Conclusions:** Selenium modulates pathogenic T-cell responses in TED, most prominently suppressing the Tfh compartment and attenuating the Tfh–B cell–autoantibody axis via ferroptosis and apoptosis. These findings suggest a mechanistic framework for the clinical benefit of selenium in mild TED and highlight the importance of dose selection within its narrow therapeutic window.

## Introduction

Thyroid Eye Disease (TED), also known as Graves’ orbitopathy, is an autoimmune inflammatory disorder of the orbit closely associated with Graves’ disease (GD). Population-based data indicate that the condition affects approximately 25–50% of patients with Graves’ hyperthyroidism, with a pooled global prevalence of around 40% (Chin et al., 2020; Bartalena et al., 2025). The annual incidence is markedly higher in women than in men (16 per 100,000 vs. 2.9 per 100,000), and disease severity increases with older age and with smoking (Bartley, 1994; Perros et al., 1993). Although the majority of patients present with mild disease—characterised by ocular irritation, conjunctival redness, and lid retraction—TED occurs in moderate-to-severe form in approximately 22% of cases, and sight-threatening complications arise in roughly 1% (Bartalena et al., 2025). Progressive disease is characterised by inflammation and expansion of the extraocular muscles and orbital adipose tissue, leading to proptosis, diplopia, exposure keratopathy, and, in severe cases, compressive optic neuropathy (Wiersinga et al., 2021).

The pathogenesis of TED is closely linked to Thyroid-Stimulating Immunoglobulins (TSI), autoantibodies directed against the Thyroid-Stimulating Hormone Receptor (TSHR). Binding TSI to TSHR results in constitutive receptor activation and unregulated thyroid hormone production. Critically, TSHR is also expressed on orbital fibroblasts and preadipocytes, and its activation by stimulatory TSHR antibodies promotes adipogenesis, extracellular matrix expansion, and glycosaminoglycan (GAG) accumulation, leading to orbital tissue enlargement and inflammation (Kumar et al., 2011; Bahn et al., 2010; Görtz et al., 2016). The insulin-like growth factor-1 receptor (IGF-1R), which is co-expressed on orbital fibroblasts and synergises with TSHR signalling, represents a further important pathogenic axis; blockade of this receptor with teprotumumab underpins one of the most significant advances in TED therapy (van Steensel et al., 2015; Bisnauthsing et al., 2026). Despite advances in targeted therapies, a subset of patients remains refractory to current treatment strategies (Bahn, 2010; Douglas et al., 2020), highlighting the need for a deeper mechanistic understanding of TED.

T cells play a central role in TED pathogenesis through both systemic immune dysregulation and local orbital infiltration. In active disease, CD4^+^ T-helper (Th) cells—predominantly of the Th1 subtype—predominate in orbital infiltrates, alongside smaller populations of Th2, Th17, and regulatory T cells (Wakelkamp et al., 2000; Fang et al., 2019). Th1-associated cytokines, particularly interferon-γ (IFN-γ) and tumour necrosis factor-α (TNF-α), are elevated in active TED and directly stimulate orbital fibroblast proliferation, hyaluronic acid synthesis, and GAG production (Smith et al., 1998; Yang et al., 2015; Bahn, 2010). As disease progresses to a more quiescent phase, a shift toward Th2-associated cytokines (IL-4, IL-10) is observed, which sustains GAG production while modulating prostaglandin synthesis (Wiersinga, 2007). Beyond Th1 and Th2 subsets, emerging evidence implicates follicular helper T (Tfh) cells—a specialised CD4^+^ subset essential for B-cell maturation and high-affinity antibody production—in the maintenance of TSI. Circulating Tfh frequencies are elevated in Graves’ disease and correlate positively with TRAb titres and thyroid hormone levels (Li et al., 2015; Zhu et al., 2019; Crotty, 2011). Their potential contribution to sustained TSI generation and orbital pathology in TED, however, remains incompletely characterised (Fan et al., 2021). The specific contribution of individual T-cell subsets to orbital pathology in TED thus remains an important, incompletely resolved question.

Selenium has gained increasing attention as a potential immunomodulatory agent in TED. From a clinical perspective, patients with TED exhibit lower circulating selenium levels than selenium-replete controls, and relative selenium insufficiency has been proposed as an independent risk factor for disease development (Khong et al., 2014; Dehina et al., 2016; Fatourechi & Heshmati, 2024). The landmark EUGOGO randomised, double-blind, placebo-controlled trial demonstrated that six months of sodium selenite supplementation (100 μg twice daily) in patients with mild GO significantly improved composite eye outcomes and quality of life, and reduced the risk of disease progression; these benefits persisted through the six-month post-treatment follow-up period (Marcocci et al., 2011). On the basis of this evidence, the European Group on Graves’ Orbitopathy (EUGOGO) incorporated selenium into its clinical practice guidelines for mild GO (Bartalena et al., 2021). At an experimental level, selenium modulates multiple aspects of adaptive immunity. In models of experimental autoimmune thyroiditis (EAT), selenium supplementation reduces Th1-, Th2-, and Th17-associated cytokine production (Li et al., 2023). Separately, selenium has been shown to reduce production of the inflammatory chemokines CXCL10, IL-23, and CCL2 in autoimmune thyroid disease contexts (Pilli et al., 2022; Zake et al., 2021). These data suggest that selenium may regulate adaptive immune responses through mechanisms extending well beyond its established antioxidant properties.

Despite the clinical evidence supporting selenium supplementation in mild TED, the immunological mechanisms that underlie its therapeutic effect remain poorly defined. In particular, no biological studies have systematically examined how selenium directly influences pathogenic T-cell subsets including Th1, Tfh, that are central to orbital inflammation and TSI generation in TED. The relative contributions of these subsets to orbital pathology, and the extent to which selenium modulates them, have not been characterised in a TED animal model.

Therefore, this study aims to investigate the effects of selenium on T-cell-mediated immune responses in a TED animal model. We hypothesise that selenium supplementation modulates pathogenic CD4^+^ T-cell subsets,specifically reducing Th1 and Tfh responses, thereby attenuating orbital inflammatory activity.

## Method

### TED Induction in Animals

A total of 122 female BALB/c mice (6–8 weeks of age) were obtained from the Laboratory Animal Services Centre of The Chinese University of Hong Kong. All animal procedures were approved by the Animal Experimentation Ethics Committee (AEEC) of The Chinese University of Hong Kong (Approval No. 22-136-MIS) and conducted in accordance with institutional guidelines.

Mice were randomly assigned to four experimental groups (n = 5 per group per experiment). The study was conducted in multiple independent experimental batches to yield a total of 122 animals. Mice were immunized with either the eukaryotic expression plasmid pTriEx1.1Neo–human TSHR A-subunit or the control plasmid pTriEx1.1Neo–β-galactosidase (β-Gal), following established protocols (Moshkelgosha et al., 2018; Philipp et al., 2022). The plasmids were generously provided by Prof. Anja Eckstein (University of Duisburg-Essen, Germany).

### Serological Analysis

Serum levels of total thyroxine (T4) and anti-thyrotropin receptor antibodies (TRAb) were quantified using commercially available enzyme-linked immunosorbent assay (ELISA) kits (MyBioSource, San Diego, CA, USA) according to the manufacturer’s instructions.

### Histopathology and Immunohistochemistry of Orbital Tissues

Orbital tissues were harvested from mice and fixed in 10% neutral-buffered formalin. To ensure complete decalcification of the orbital bones, tissues were soaked in Osteosoft™ solution (Sigma-Aldrich, USA) for 48 hours at room temperature. Sections were mounted on glass slides and stained with Haematoxylin and Eosin to visualise adipose tissue composition, including brown and white fat. In addition, Masson’s trichrome staining was performed to assess the extent of collagen deposition and fibrosis within the orbital tissue.

### Immunohistochemistry

Immunohistochemical staining was performed to detect T-cell infiltration in orbital tissues. Sections were incubated overnight at 4°C in a humidified chamber with primary antibodies against CD3 (1:100) and CD4 (1:100), diluted in antibody diluent. After washing in PBS, slides were incubated with species-appropriate HRP-conjugated or fluorophore-conjugated secondary antibodies for 1 hours at room temperature.

For chromogenic detection, sections were developed using 3,3′-diaminobenzidine (DAB) substrate and counterstained as appropriate. For fluorescent detection, sections were mounted using antifade mounting medium containing DAPI without dehydration or xylene clearing.

Stained sections were imaged using either a brightfield or fluorescence microscope

### Flow Cytometry Analysis

Spleens were harvested from BALB/c mice 8 weeks post-immunization and were mechanically ground with a 40 µm cell strainer to generate a single-cell suspension. The suspension was washed with 5 mL PBS and centrifuged at 400 × g for 3 minutes at 4 °C. The red blood cell was lysed with RBC lysis buffer and the splenocyte pellet was resuspended in complete RPMI 1640 medium containing 10% FBS. The isolated splenocytes were then stained APC-R700 IFNγ, BV605 CD4⁺(BD Biosciences, US), APC IFNγ, IL-17A PE, APC CXCR5, PE-Cyanine 7 PD-, CD-19 (BioLegend).

Isolated splenocytes were stimulated with 50 mg/mL phorbol 12-myristate-13-acetate (PMA) and 750 mg/mL ionomycin for 4 hours at 37 °C in the presence of 5 µg/mL brefeldin A to inhibit cytokine secretion. Following stimulation, cells were fixed with 4% paraformaldehyde and incubated for 30 minutes at room temperature. Cells were then permeabilized and stained with fluorescently labelled antibodies against intracellular markers such as IFN-γ and IL-17A for 30 minutes at room temperature. Flow cytometric analysis was performed using a BD FACSLyric Flow Cytometry System (12-colour capability).

### Isolation of Tfh Cells and Selenium Treatment in vitro

CD4⁺ T cells were isolated from mouse spleens using the EasySep™ Mouse CD4^+^ T Cell Isolation Kit (Stem Cell Technologies, Canada) according to the manufacturer’s instructions. Cells were seeded in 24-well plates at a density of 500,000 cells per well in 1 mL of RPMI supplemented with 10% fetal bovine serum (FBS) and 1% penicillin/streptomycin (P/S). Sodium selenite (Sigma-Aldrich, USA) was applied to the cells at concentrations of 0, 1 µM, 5 µM, 10 µM, 20 µM, and 50 µM for 24 hours.

To assess cell death mechanisms, apoptosis was measured using Annexin V-FITC staining (Thermo Fisher Scientific, USA), and lipid peroxidation, a marker of ferroptosis, was detected using BODIPY 581/591 C11 dye (Thermo Fisher Scientific, USA), followed by flow cytometric analysis.

### Selenium Treatment of TED Mice

Subsequently, mice were fed a low-selenium diet (0.07 ppm Se; Labdiet, USA) *ad libitum* for one week prior to plasmid immunisation, in contrast to the regular diet containing 0.41 ppm selenium. They continued on the low-selenium diet until the experiment’s conclusion, when they were sacrificed. Sodium selenite supplementation at 0.2 mg/kg was administered daily for 10 days following the final immunization with prior experiment for determine the toxicity of selenium (Larrouquère et al., 2021).

### IL-21 Measurement in vitro and in vivo

IL-21 levels were measured using an ELISA kit (Invitrogen, Thermo Fisher, USA) according to the manufacturer’s instructions. Serum samples were collected from four groups of mice: β-gal immunisation without selenium (Se) treatment, β-gal with Se treatment, TSHR immunisation without Se, and TSHR with Se treatment. Additionally, IL-21 was quantified as the culture supernatants of Se-treated Tfh cells. For in vitro analysis, CD4⁺ T cells were isolated from spleens using the EasySep™ Mouse CD4^+^ T Cell Isolation Kit (Stemcell Technologies, Canada). Tfh cells were further purified by labelling with APC-conjugated CXCR5 antibody, followed by isolation using the EasySep™ APC Positive Selection Kit II (Stemcell). Purified Tfh or CD4⁺ cells were cultured in 96-well plates for 24 hours with sodium selenite at concentrations of 1 µM, 5 µM, 10 µM, and 20 µM. After incubation, cells were collected for flow cytometry, and culture supernatants were harvested for IL-21 quantification by ELISA (Thermo Fisher, USA).

### Statistics and power calculation

All statistical analyses were performed using GraphPad Prism 7. Data are presented as mean ± standard error of the mean (SEM). Comparisons between groups were conducted using the Student’s t-test. Statistical significance was denoted as follows: *p* < 0.05 (\**), p < 0.01 (**), p* < 0.001 (***), and *p* < 0.0001 (****). The power calculation of sample size was performed a priori using G*Power and confirmed analytically. For pairwise two-sample comparisons (two-tailed t-test, α = 0.05, power = 0.80.

## Results

### Selenium Dosage Evaluation

To determine the therapeutic effect of selenium in the experimental TED animal model, the non-toxic dosage range of selenium in wild-type mice. Based on previous research by Larrouquère et al. (2021), which indicated that doses up to 2.25 mg/kg exhibit minimal toxicity in mice, this study evaluated the safety and tolerability of orally administered selenium across a dose range of 0.1–4 mg/kg/day for 10 consecutive days in female BALB/c mice. Throughout the treatment, the body weights of the mice were recorded daily as a general indicator of health and toxicity. Our results demonstrated that doses at or below 0.2 mg/kg had no significant adverse effect on body weight, and all mice survived without observable clinical signs of toxicity (Figure 1). In contrast, higher doses above this threshold showed signs of weight loss and increased morbidity, suggesting potential toxicity. Consequently, 0.2 mg/kg was selected as the optimal, non-toxic dose for subsequent selenium treatment experiments in the TED mouse model.

**Figure 1:**
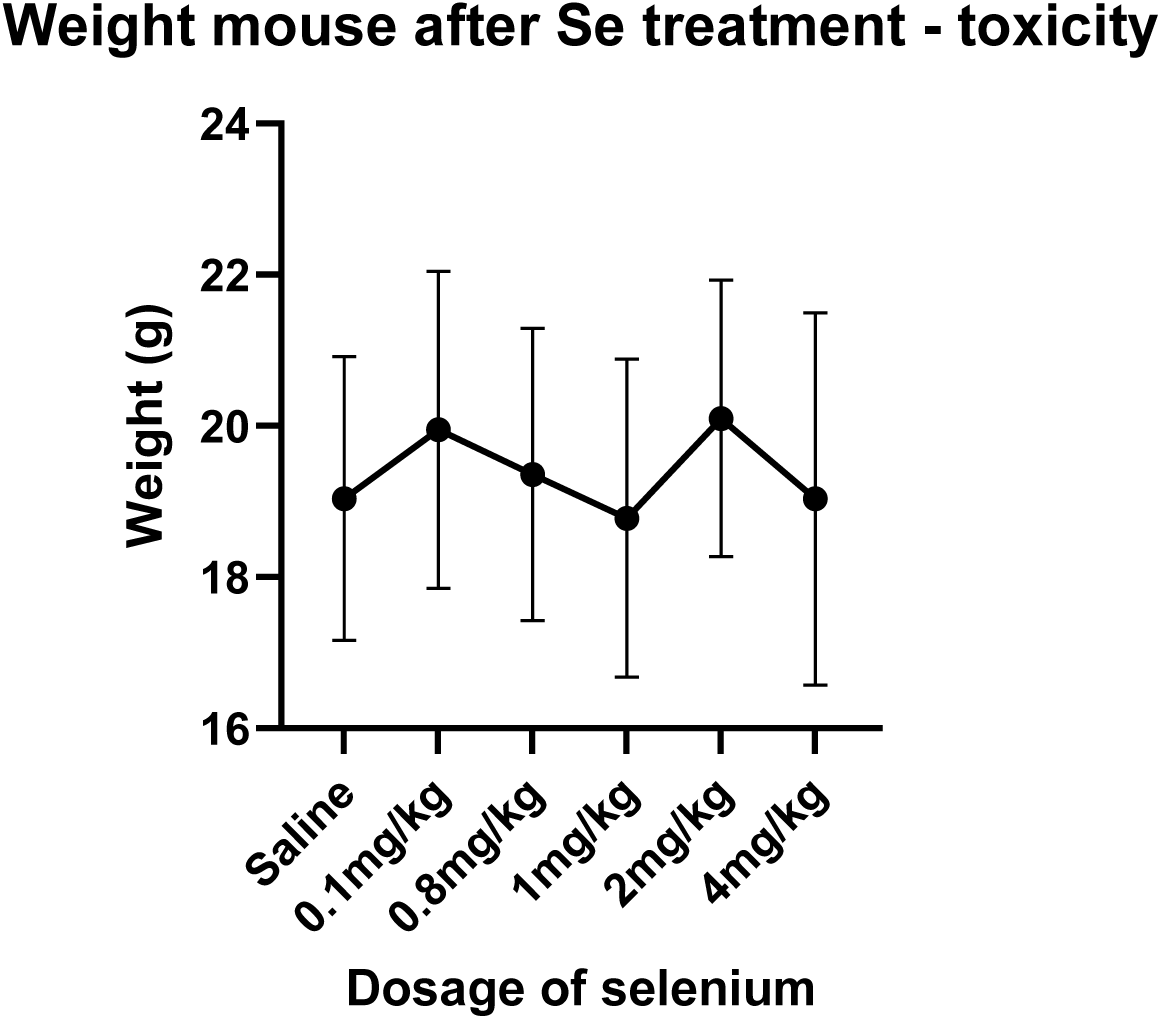
Assessment of selenium toxicity and its effects on body weight in mice. Mice (n = 4 per group) were administered selenium at doses ranging from 0.1 mg/kg to 4 mg/kg body weight. Within the range of 0.1 –0.8 mg/kg, no signs of selenium overdose or adverse effects on body weight were observed. Data are presented as mean ± SEM.

### Immunohistochemistry of CD3⁺ T Cells, Masson-Trichome and H&E Staining of orbital Tissues

Compared to the β-gal plasmid-immunised control group, mice immunised with the TSHR plasmid exhibited a significant increase in CD3^+^ T cell infiltration within the orbital tissues, confirming an enhanced immune response characteristic of TED pathology. Notably, daily administration of selenium at 0.2 mg/kg for 10 consecutive days led to a decrease in CD3⁺ cell infiltration in these tissues, indicating a potential immunomodulatory effect of selenium treatment (Figure 2 and 3).

**Figure 2:**
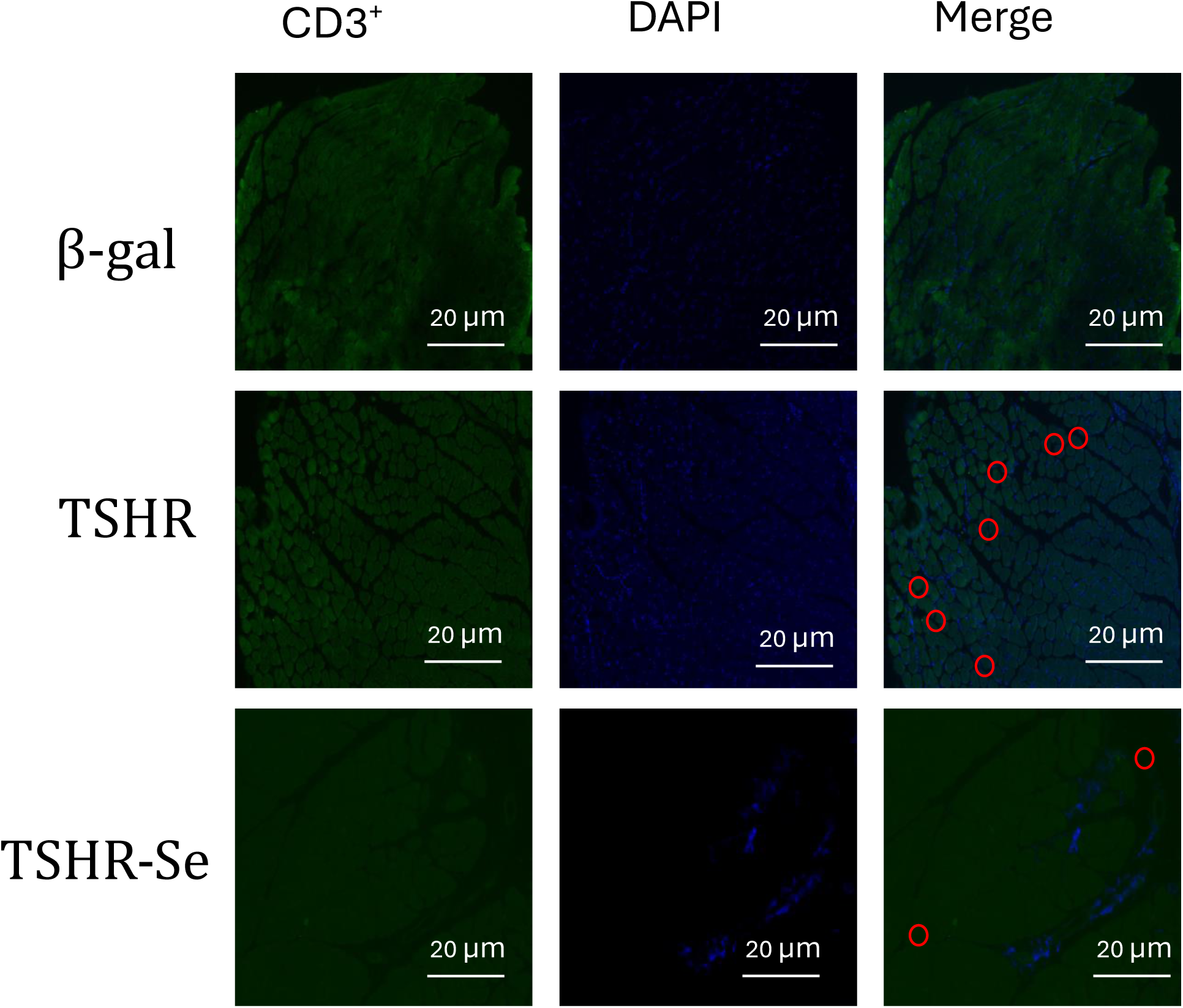
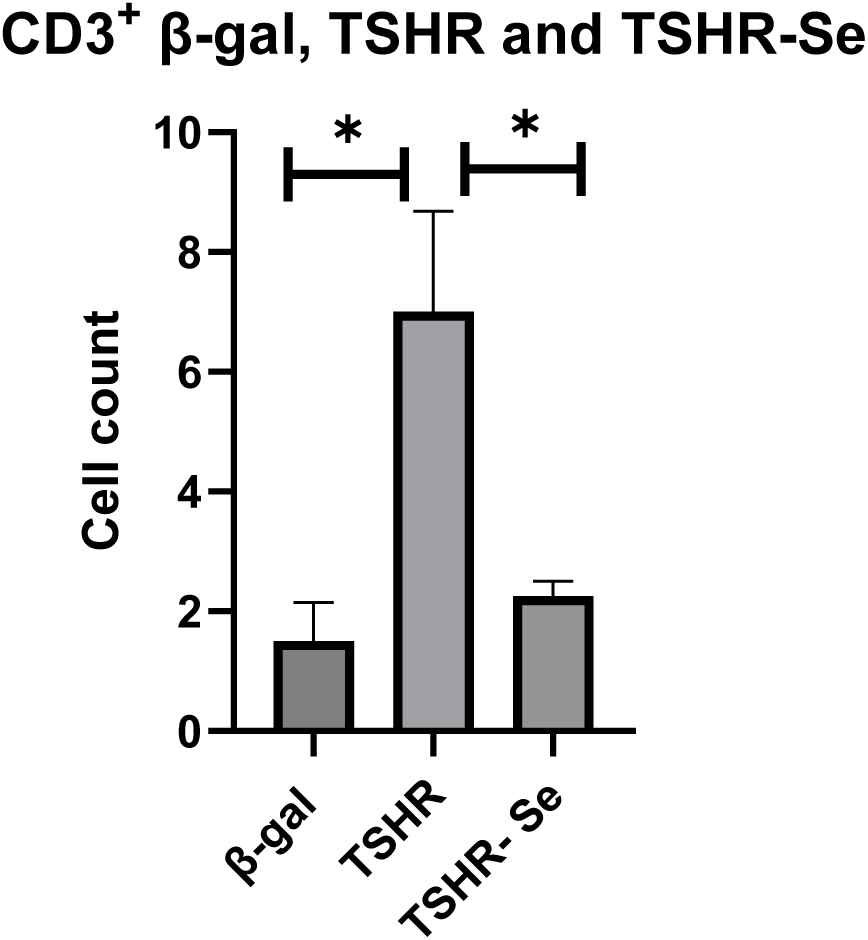
Immunohistochemical staining of CD3⁺ T cells in orbital tissues. Sections were stained using a rabbit anti-mouse CD3 antibody. Images were captured at 40× magnification, highlighting the distribution and infiltration of CD3⁺ cells within the orbital tissue. Scale bar = 20 µm.

**Figure 3:**
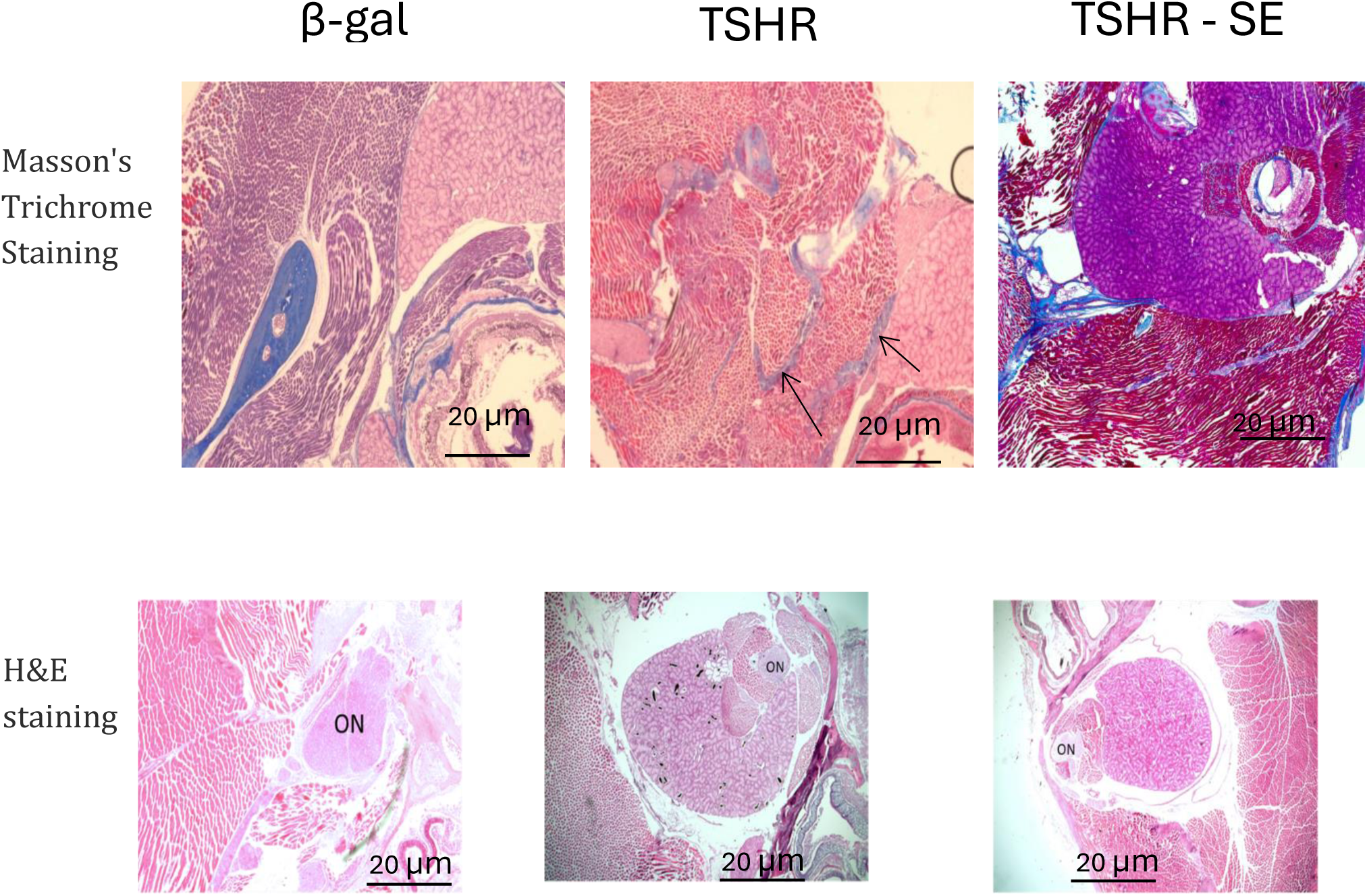
Masson’s Trichrome and Haematoxylin and Eosin (H&E) staining of orbital tissues. Arrows indicate areas of collagen fibrosis highlighted by Masson’s Trichrome staining. Anatomical landmarks including the optic nerve (ON) and Harderian gland (HG) are labelled. Images were captured at 40× magnification. Scale bar = 20 µm.

### Serological Studies

Serological analysis revealed that TSHR plasmid-immunised mice exhibited significantly elevated serum levels of total thyroxine (T4) and thyrotropin receptor antibodies (TRAb) compared to β-gal plasmid-immunised controls, confirming the successful induction of autoimmune thyroid activity (Figure 4). Notably, treatment with selenium at 0.2 mg/kg in the TSHR-immunised group resulted in a significant reduction in both T4 and TRAb levels, indicating that selenium administration may modulate thyroid hormone dysregulation and autoantibody production in this experimental model of TED (Figure 4).

**Figure 4:**
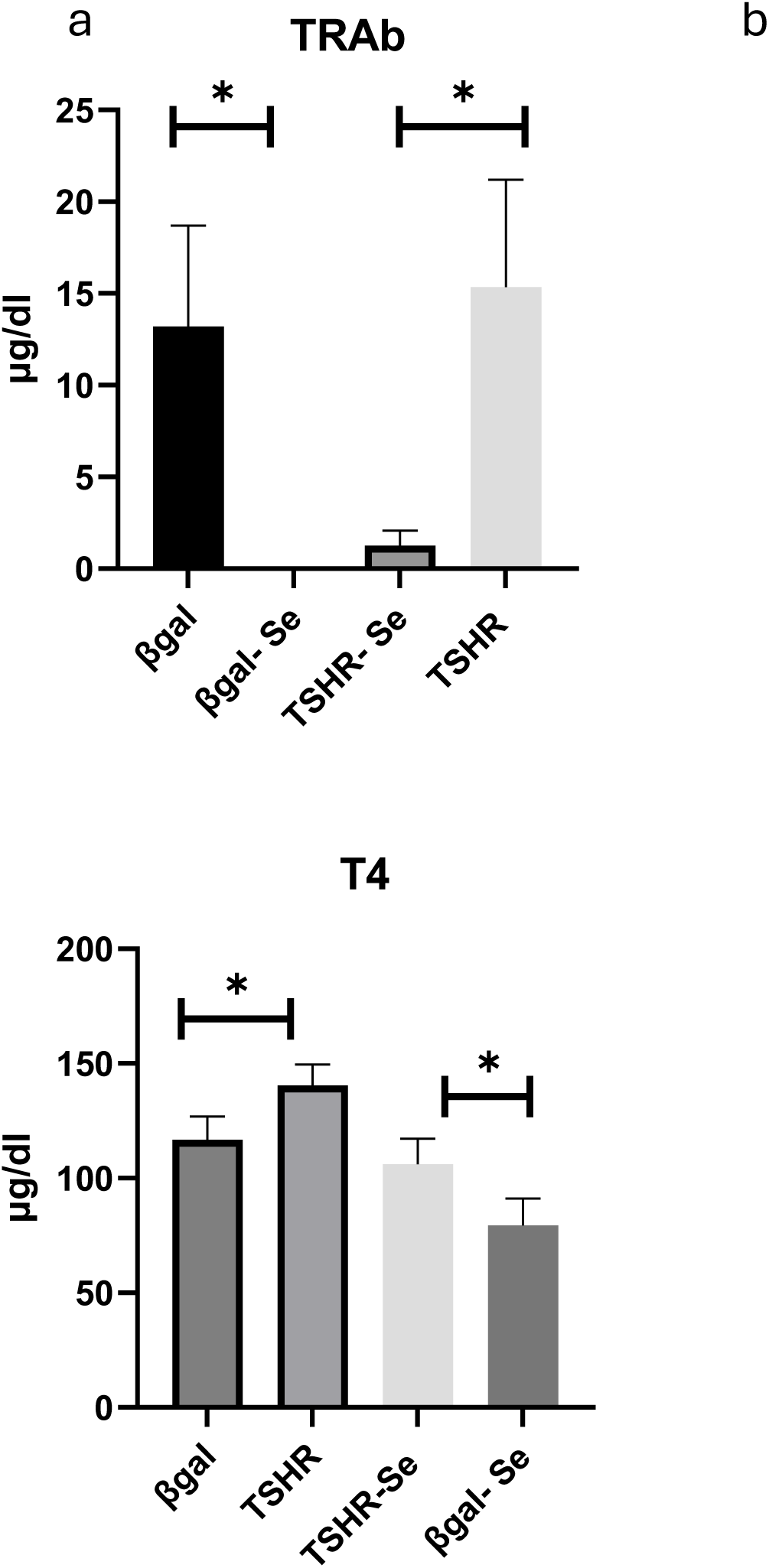
Effects of selenium treatment on serum T4 and TRAb levels in TSHR- and β-gal immunised mice. Serological measurements of total thyroxine (T4, b) and thyroid-stimulating hormone receptor antibodies (TRAb, a) in mice immunised with TSHR or β-gal plasmids, with or without selenium (Se) treatment. Significant differences between groups are indicated by * (p < 0.05). Sample size: n = 16 per group. Data are presented as mean ± SEM and analysed using an unpaired Student’s t -test.

### T Cell Subtype Abundance in TED mice Treated With Selenium

Considering that selenium present in the regular mouse diet could influence the effects of additional selenium supplementation, we optimised our experimental design by feeding mice a low-selenium diet (0.07 ppm Se) starting one week prior to plasmid immunisation and continuing until sacrifice. Throughout this period, mice received supplemental selenium at 0.2 mg/kg. Following treatment, spleens were harvested for T cell isolation, staining, and flow cytometry analysis. Our findings demonstrated a significant reduction in the abundance of Th1 and T follicular helper (Tfh) cells in TSHR plasmid-immunised mice receiving selenium supplementation, while Th17 cell levels remained unchanged (Figure 5).

**Figure 5:**
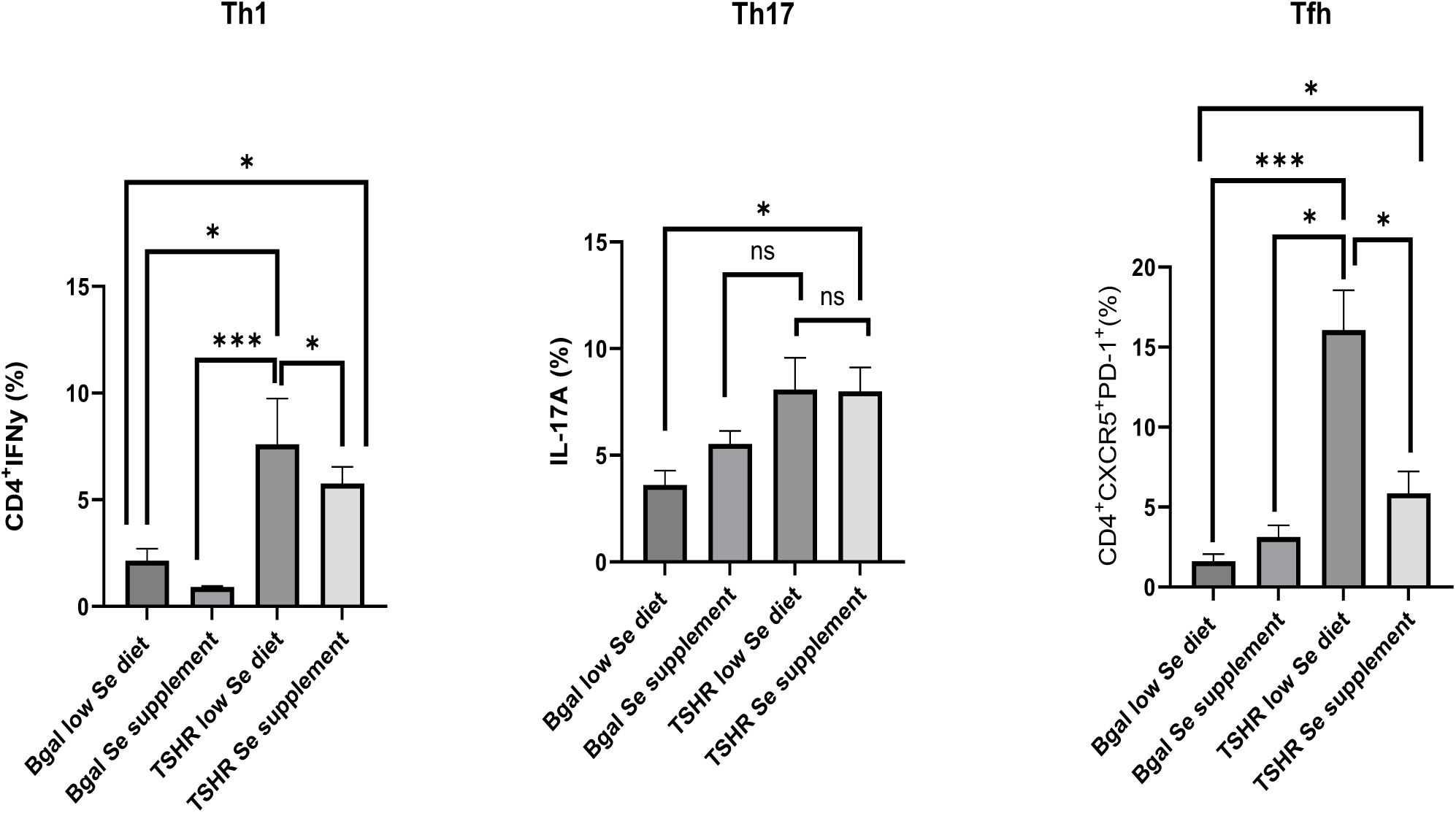
Effects of Selenium Supplementation on Th1, Th17, and Tfh Cell Abundance in TED Mice. Abundance of Th1, Th17, and Tfh cells in thyroid eye disease (TED) mice fed a low-selenium diet and subsequently treated with additional selenium supplementation (n = 10 –17 per group). selenium treatment significantly modulated T cell subset abundance compared to untreated controls. *p < 0.05, ***p < 0.001; ns indicates no statistical significance. Data are presented as mean ± SEM and analysed using an unpaired Student’s t test.

To validate these observations, a parallel experiment was conducted using mice maintained on a regular selenium-containing diet (0.44 ppm Se). In TSHR plasmid-immunised mice, those fed a low-selenium diet exhibited elevated levels of Tfh, Th1, and Th17 cells compared to mice on the regular diet (Figure 5). Moreover, supplementation with selenium in mice on the low-selenium diet resulted in a significant decrease in the abundance of Tfh, Th1, and Th17 cells (Figure 6).

**Figure 6:**
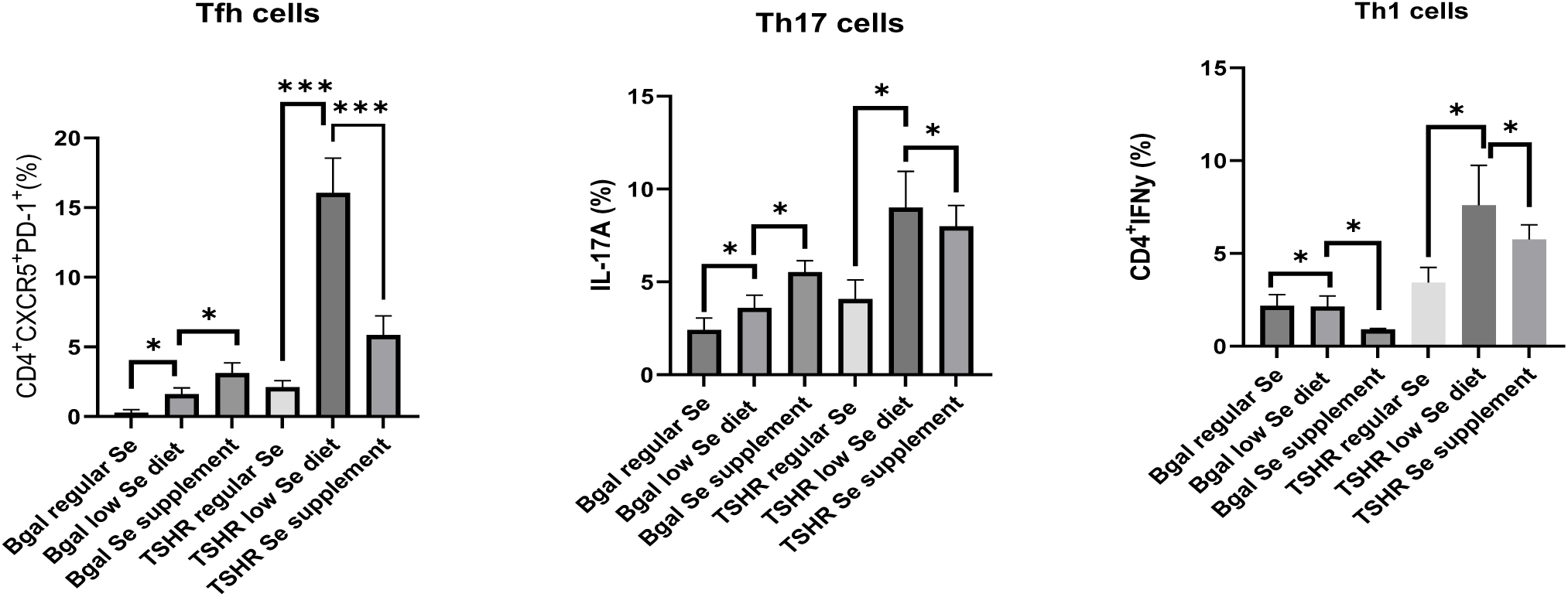
Mice were immunised with β-gal and TSHR plasmids and fed either a regular Se diet or a low-Se diet. An additional group of mice was first fed a low-Se diet and then received Se supplementation. Sample sizes ranged from n = 9 to n = 26 per group. Spleens were collected for T cell isolation, staining, and flow cytometry analysis to assess the abundance of Tfh, Th1, and Th17 cells. ns indicates no statistical significance. * and *** denote p <0 .05 and p <0 .001, respectively.

### IL-21 Concentration

Tfh cells play a pivotal role in humoral immunity by secreting interleukin-21 (IL-21), which facilitates B cell activation and promotes antibody production. Accordingly, we measured serum IL-21 concentrations in the experimental groups (Figure 7). The results demonstrated a significant elevation of IL-21 in TSHR plasmid-immunised mice, which was markedly reduced upon 10 days of selenium supplementation (in vivo).

**Figure 7:**
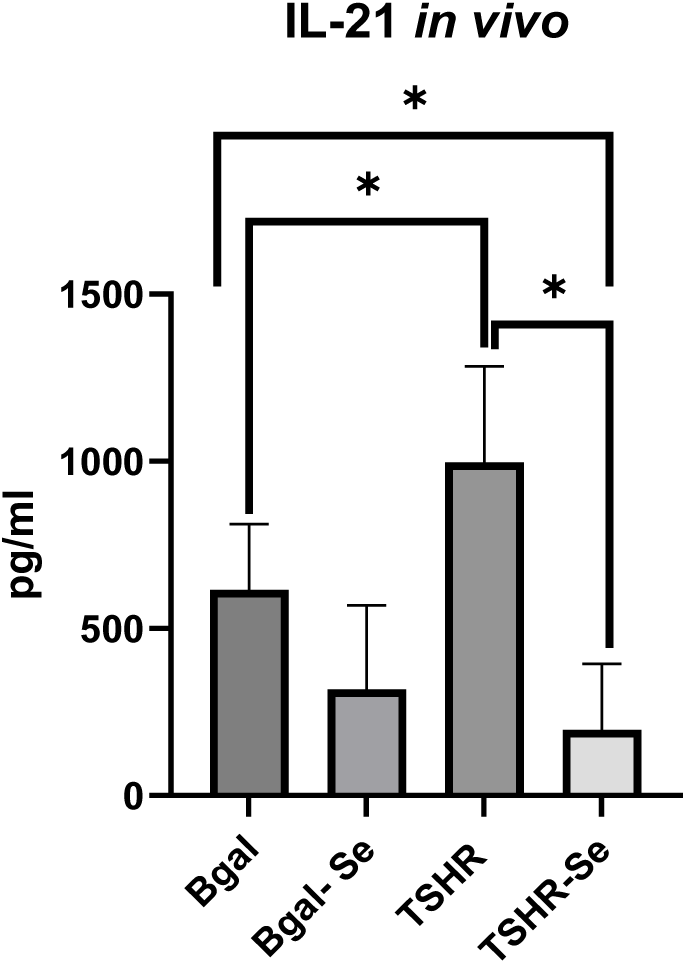
Effect of Selenium on IL-21 Expression in β-gal and TSHR-Immunised Mice. Mice were immunised with β-gal and TSHR plasmids and maintained on a selenium-supplemented diet (n = 4 per group). ELISA measured IL-21 levels. Data are presented as mean ± SEM. * indicates p <0.05.

To further investigate this finding in vitro, we isolated Tfh cells from TED mice and treated them with varying concentrations of selenium (0 to 10 µM) in vitro (Figure 8). Interestingly, a dose-dependent trend towards increased IL-21 secretion was observed, indicating that selenium may directly stimulate IL-21 production by Tfh cells.

**Figure 8:**
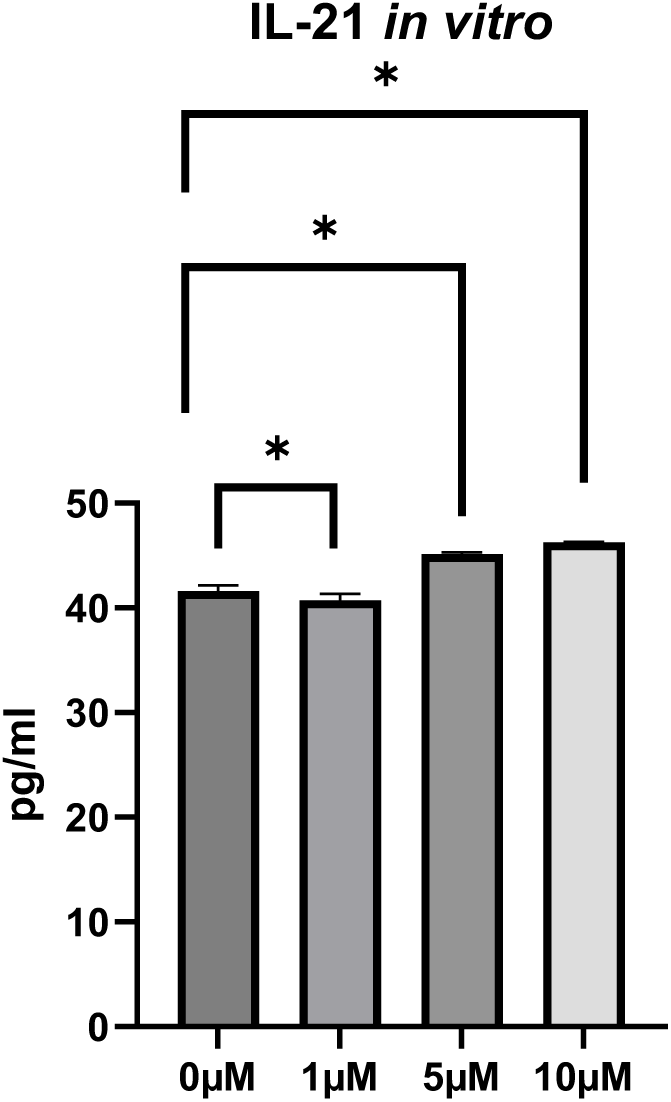
In Vitro Selenium Treatment Alters IL-21 Production by Tfh Cells. Measurement of IL-21 in vitro. Tfh cells isolated from TED mice were treated with increasing concentrations of selenium (Se) (n = 3 per group). IL-21 expression was measured. * indicates p <0.05.

Given the role of IL-21 in B cell activation, we further evaluated B cell abundance of β-gal and TSHR plasmid-immunised mice, with and without selenium supplementation (Figure 9). Consistent with disease pathology, B cells were significantly more abundant in TED mice, whereas selenium treatment markedly reduced their infiltration. This reduction in B - cells was observed following in vivo administration of selenium supplementation (Figure 9).

**Figure 9:**
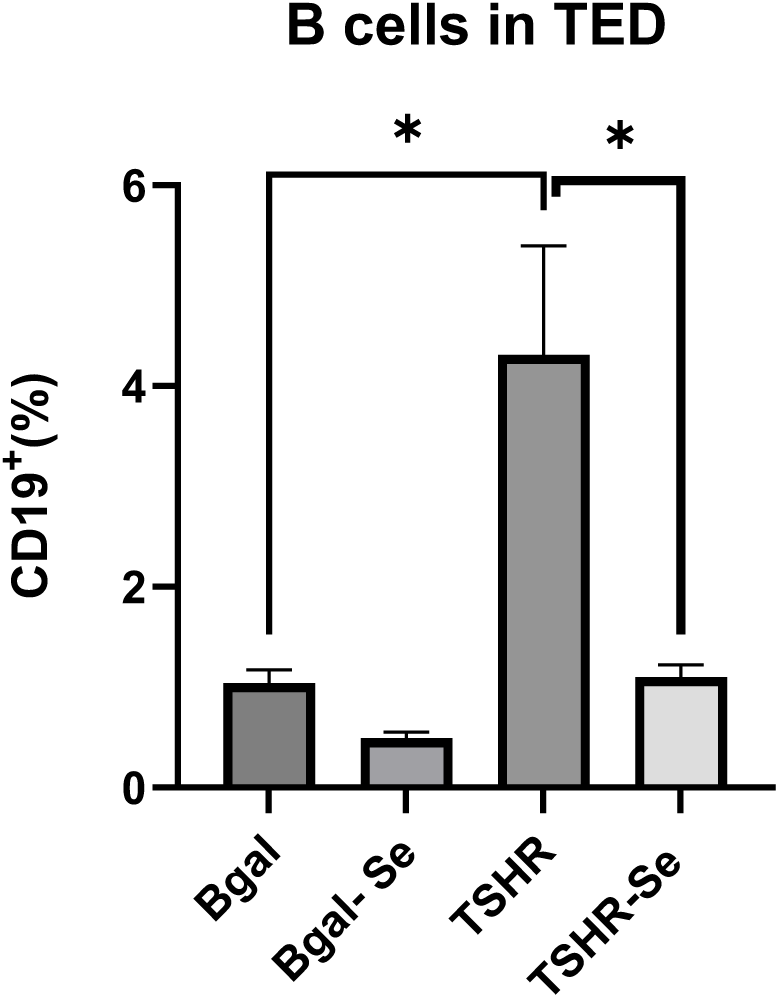
Effect of Selenium Supplementation on B Cell Abundance in Plasmid-Immunised Mice. B cell abundance was measured in mice immunised with β-gal or TSHR plasmids and fed with or without selenium (Se) supplementation (n = 9 per group). * denotes statistical significance (p <0.05).

### Cell Death in Tfh, Th1, and Th17 Cells

The observed modulation in T cell subset abundance following selenium treatment may be attributable to Se-induced alterations in cell death pathways. To investigate this, T cells were isolated from TED mice and exposed to a gradient of selenium concentrations (0, 1, 5, 10, and 20 µM) in vitro for 24 hours. Subsequently, these cells were stained with markers specific for Tfh, Th1, and Th17 subsets and analysed via flow cytometry (Figure 10). The results revealed a dose-dependent significant reduction in Tfh cell abundance caused by increased Se concentrations, accompanied by marked elevations in both apoptosis and ferroptosis. In Th1 cells, although a trend towards reduced cell numbers and heightened apoptosis was observed with higher Se levels, these changes did not achieve statistical significance; however, ferroptosis was significantly augmented in response to selenium. Conversely, Th17 cells displayed no significant alterations in abundance, apoptosis, or ferroptosis across the tested Se concentrations. These findings suggest that selenium selectively influences T cell viability and death mechanisms, particularly affecting Tfh and Th1 populations, which may contribute to the immunomodulatory effects observed in TED. Taken together, these in vitro findings demonstrate a dose-dependent biphasic response to selenium: at lower concentrations (1 µM), Tfh cell viability was maintained and IL-21 production was preserved, whereas at higher concentrations (10–20 µM), both apoptosis and ferroptosis were significantly elevated alongside increased IL-21, consistent with a cellular stress response preceding cell death. These data suggest that the immunomodulatory benefit of selenium is confined to a narrow concentration window, below which cells remain functional and above which cytotoxic mechanisms predominate.

**Figure 10:**
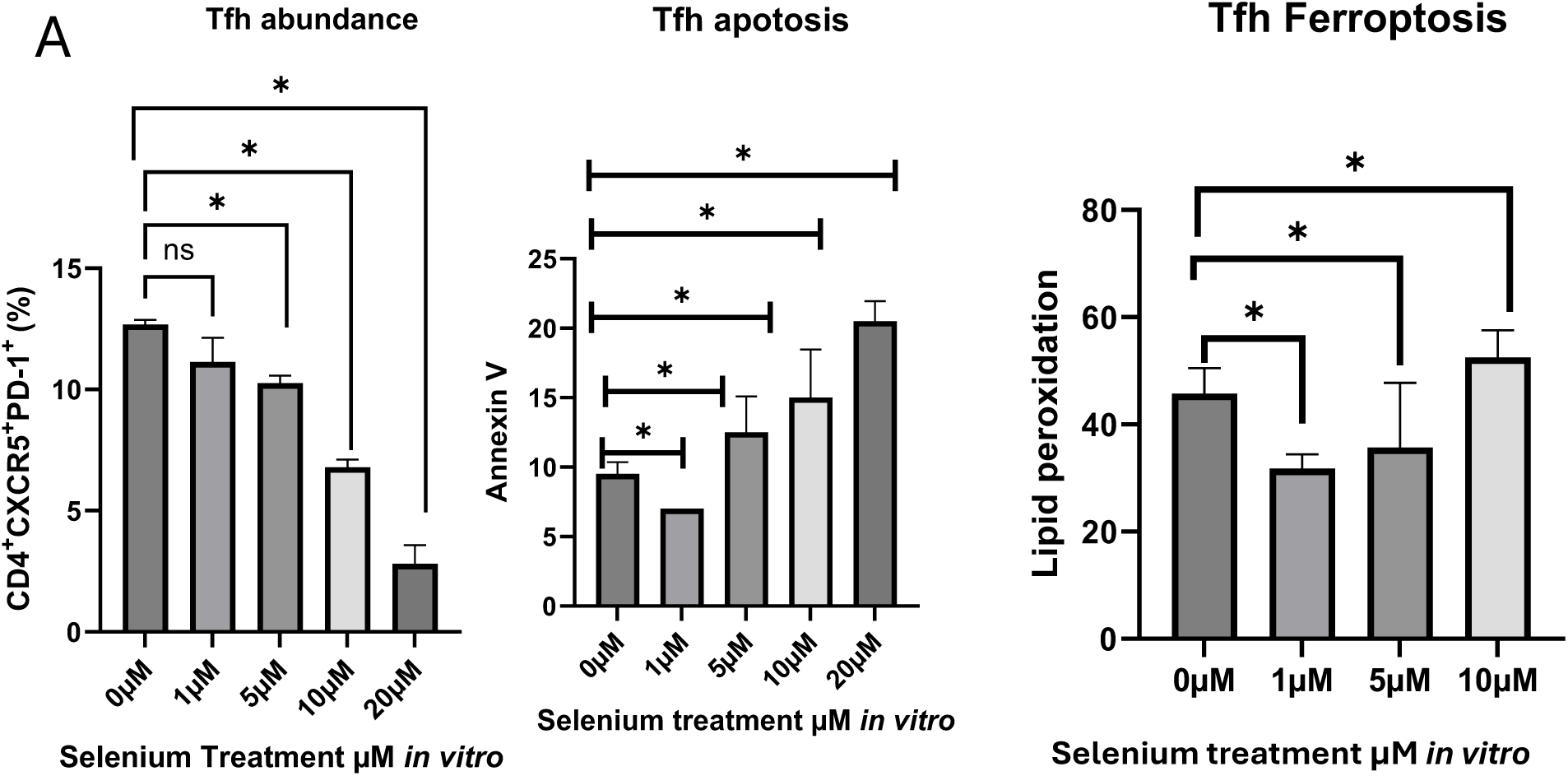
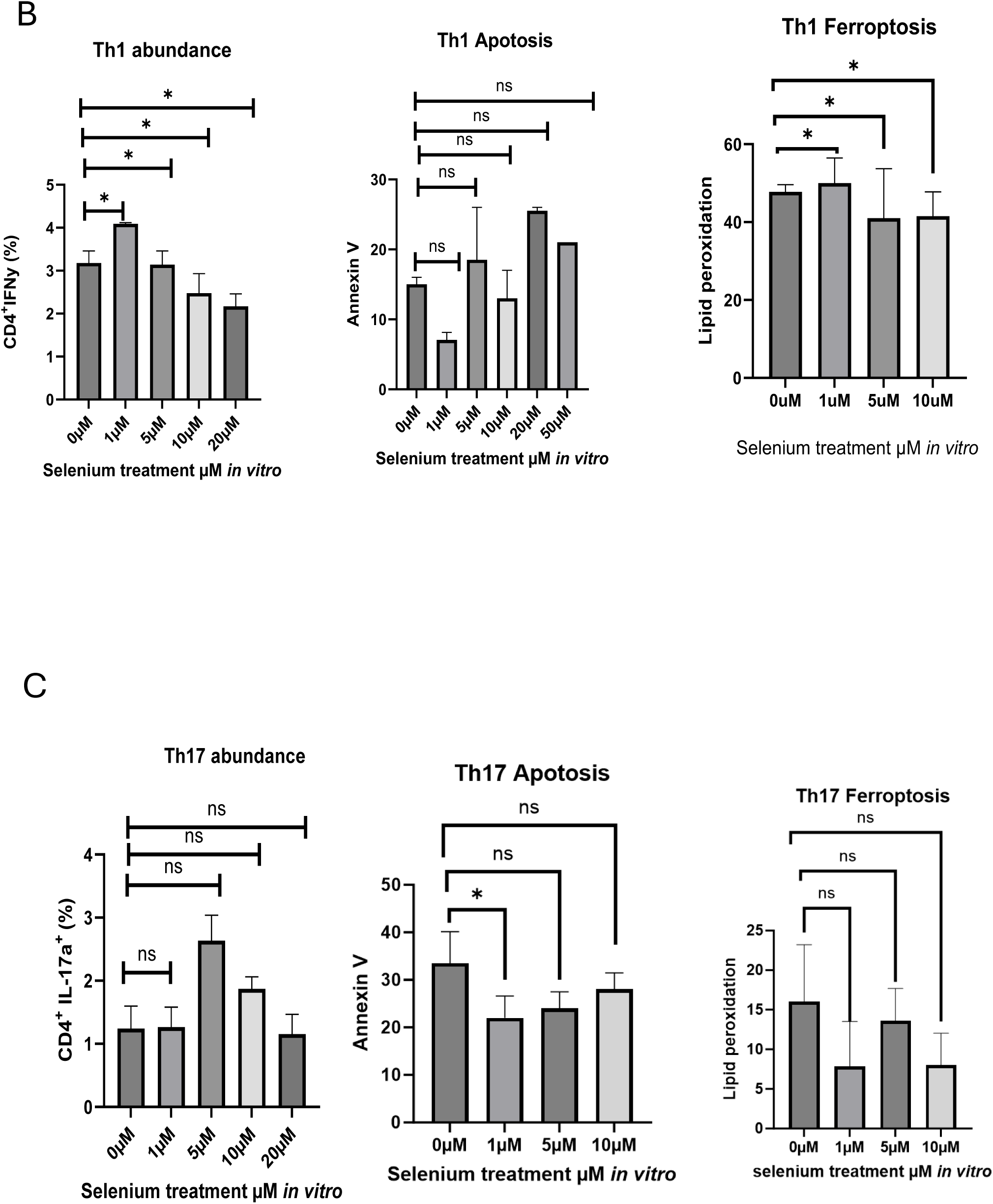
Selenium Treatment Alters Tfh, Th1, and Th17 Cell Abundance, Apoptosis, and Ferroptosis. Panels (A–C) show the effects of selenium (Se) treatment on the abundance, apoptosis, and ferroptosis of Tfh (A), Th1 (B), and Th17 (C) cells isolated from immunised mice. Data represent mean ± SEM from n = 4 mice per group. Statistical significance was determined using a Student’s t-test. ns indicates no statistical significance; * denotes p <0.05.

## Discussion

The present study investigated the immunomodulatory effects of selenium (Se) in a murine model of Thyroid Eye Disease (TED), with a particular focus on Se’s influence on pathogenic T-cell subsets, orbital inflammation. Our findings are consistent with the clinical evidence demonstrating a beneficial role for Se in mild Graves’ orbitopathy. The landmark EUGOGO randomised, double-blind, placebo-controlled trial (Marcocci et al., 2011) established that six months of sodium selenite supplementation (100 μg twice daily) significantly improved quality of life and clinical activity scores, and slowed disease progression in patients with mild TED. The mechanistic basis for this benefit, however, has remained poorly understood, and our data provide novel insight into the cellular and molecular pathways through which Se exerts its effects. Se is well established to reduce oxidative stress, suppress proinflammatory cytokines, lower thyroid autoantibodies, and modulate immune responses (Rotondo Dottore et al., 2017; Pilli et al., 2015). It is also recognized that Se exhibits a narrow therapeutic window, with toxicity emerging at supra-nutritional doses (Hadrup & Ravn-Haren, 2023).

In vivo, immunohistochemistry and Masson’s Trichrome staining demonstrated that Se treatment initiated after the second TSHR plasmid injection reduced CD3⁺ T-cell infiltration into orbital tissues and attenuated collagen fibrosis, consistent with a partial resolution of the orbital inflammatory phenotype. These findings align with prior work by Chang et al. (2010) in a comparable murine TED model. Notably, however, Se did not completely reverse the pathological features of TED, indicating that Se alone has limited standalone efficacy and is unlikely to replace established immunosuppressive or biological therapies. This is consistent with the clinical experience in which Se benefits are most pronounced in mild disease and do not extend to moderate-to-severe or sight-threatening TED (Bartalena et al., 2021). This partial efficacy may reflect the narrow therapeutic window of Se, the short treatment duration in murine models, or the inherent species-specific differences in orbital fibroblast biology that must be considered when extrapolating findings to human disease.

To further delineate Se’s immunological effects, we compared T-cell populations in mice fed a standard diet (0.41 ppm Se) versus those on a low-Se diet (0.07 ppm Se) that were subsequently supplemented with 0.2 mg/kg Se. Se-deficient conditions were associated with a significant expansion of T-cell populations, consistent with the hypothesis that Se insufficiency promotes immune hyperactivation. Upon Se supplementation, T-cell numbers declined, with the most pronounced reduction observed in follicular helper T (Tfh) cells, exceeding reductions in Th1 or Th17 populations. Tfh cells are a specialised CD4⁺ subset critical for germinal centre reactions, B-cell affinity maturation, and high-affinity antibody production, including TSI (Crotty, 2011). These data suggest that Se may partially attenuate autoantibody generation by suppressing the Tfh compartment, complementing prior observations of elevated Tfh frequencies in Graves’ disease that correlate positively with TRAb titres (Zhu et al., 2019). Se reduced Th1 and Th17 populations, consistent with prior reports in experimental autoimmune thyroiditis (EAT) models (Li et al., 2023), and with the known role of IFN-γ and IL-17 in activating orbital fibroblasts and amplifying hyaluronic acid synthesis in TED (Fang et al., 2019).

The reduction in Tfh cells observed here appears to contradict the findings of Yao et al. (2021), who demonstrated in a non-autoimmune context that physiological Se supplementation protects Tfh cells from ferroptosis via GPX4 upregulation, thereby enhancing germinal centre responses and antibody titres following influenza vaccination. This apparent discrepancy is most likely explained by context- and dose-dependent differences in Se’s effects. In the immunologically activated, Se-deficient setting of TED, where Tfh cells are already aberrantly expanded, Se supplementation may restore redox homeostasis by selectively restraining pathogenic Tfh activity, rather than uniformly protecting all Tfh cells from ferroptosis as in a vaccination context. Moreover, the doses employed in our study differ from those in Yao et al. (2021), and the autoimmune versus non-autoimmune background of the respective models likely shapes the immunological outcome. The response of Tfh cells to Se thus appears highly context-dependent, modulated by baseline selenium status, the inflammatory milieu, and the dose of supplementation. Our in vitro experiments, Tfh cells survived at 1 μM Se, but exposure to 10 μM Se was associated with increased IL-21 expression alongside evidence of ferroptosis and apoptosis. Ferroptosis is an iron-dependent, non-apoptotic form of programmed cell death driven by the accumulation of phospholipid hydroperoxides, regulated principally by the selenoenzyme GPX4 (Matsushita et al., 2015; Yao et al., 2021). A key finding of this study is the dose-dependent, biphasic nature of selenium’s effects on Tfh cells in vitro. At low concentrations approximating physiological levels, selenium appeared to support cell survival and was associated with maintained or modestly increased IL-21 production, consistent with the known role of GPX4-mediated antioxidant defence in sustaining T-cell function (Yao et al., 2021). In contrast, at higher concentrations, selenium induced significant ferroptosis and apoptosis in Tfh cells, accompanied by a stress-associated rise in IL-21 prior to cell death. This biphasic pattern aligns with selenium’s well-recognised narrow therapeutic window (Hadrup & Ravn-Haren, 2023) and has direct clinical relevance: the beneficial effects observed in the EUGOGO trial (Marcocci et al., 2011) and in our in vivo model are most plausibly attributable to selenium operating within the lower, cytoprotective range, where antioxidant and immunomodulatory effects predominate without inducing cellular toxicity. Exceeding this range, as modelled by our higher in vitro doses, shifts the balance toward non-selective cytotoxicity, which would be counterproductive in a clinical setting.

Low-dose selenium supplementation improves TED symptoms most plausibly through antioxidant mechanisms, whereas excessive supplementation may paradoxically impair immune function (Hadrup & Ravn-Haren, 2023). This is consistent with the observation that not all studies have demonstrated immunological benefits of selenium, particularly in patients with more active autoimmune thyroid disease, underscoring that selenium’s effects are both dose-and disease-context-dependent (Karanikas et al., 2008). At concentrations excessive reactive oxygen species generated by high selenium exposure can impair T-cell activation and survival by inducing mitochondrial dysfunction, apoptosis, and ferroptosis via the GPX4 axis (Matsushita et al., 2015; Yao et al., 2021). Taken together, these findings highlight that selenium’s immunomodulatory benefit in TED operates within a narrow therapeutic window: physiological supplementation selectively modulates pathogenic T-cell subsets and attenuates orbital inflammation, whereas supraphysiological doses shift the balance toward non-selective cytotoxicity. This dose-dependency has direct implications for both clinical practice and the design of future selenium supplementation trials in TED, where careful dose selection and patient stratification by baseline selenium status will be essential.

### Strengths

This study has several notable strengths. The use of a controlled low-selenium dietary model prior to supplementation provides a more rigorous experimental baseline than studies conducted under standard dietary conditions, allowing the immunological effects of selenium repletion to be more precisely attributed. This is complemented by a multi-level experimental design that integrates in vivo immunisation, flow cytometric immunophenotyping, serological analysis, and in vitro dose–response assays, generating internally consistent evidence across multiple biological levels. Importantly, this is among the first studies to characterise the effect of selenium specifically on the Tfh–B cell axis in a TED model, extending the mechanistic focus beyond the Th1/Th2 paradigm that has traditionally dominated the field and linking selenium’s immunomodulatory effects to autoantibody generation via IL-21 and B cell suppression. The dose–response analysis of selenium’s effects on Tfh cell viability, apoptosis, and ferroptosis further provides a plausible biological framework for understanding why low-dose supplementation may be clinically beneficial while higher doses carry toxicity risk — an observation that aligns with the narrow therapeutic window recognised in clinical practice. Taken together, the identification of ferroptosis as a potential mechanism through which selenium modulates pathogenic T-cell populations represents a novel mechanistic contribution to TED immunology that has not previously been explored in this disease context.

### Limitations

Certain limitations must be acknowledged. Se supplementation reduced, but did not fully reverse, orbital inflammation and fibrosis, highlighting its limited standalone efficacy and the narrow therapeutic window of Se—a recognised translational challenge (Hadrup & Ravn-Haren, 2023). Our immunophenotyping focused on Tfh, Th1, and Th17 subsets and did not comprehensively characterise the full orbital immune landscape, including macrophages, orbital fibroblasts, and endothelial cells, all of which contribute to TED pathogenesis. The short duration of Se treatment in murine models may not adequately reflect the long-term dynamics of a chronic human condition, and established species-specific differences in orbital anatomy and fibroblast biology must be considered when extrapolating results. Finally, the in vitro concentrations used (1–10 μM Se) exceed those typically achievable through dietary supplementation, and the translational relevance of the high-dose ferroptotic effects to clinical Se supplementation at 100–200 μg/day warrants cautious interpretation.

## Notes

### Competing Interest Statement

The authors have declared no competing interest.

